# Strain-specific evolution and host-specific regulation of transposable elements in the model plant symbiont *Rhizophagus irregularis*

**DOI:** 10.1101/2023.12.01.569630

**Authors:** Jordana Inácio Nascimento Oliveira, Nicolas Corradi

## Abstract

Transposable elements (TEs) are repetitive DNA that can create variability in genome structure and regulation. The genome of *Rhizophagus irregularis*, a widely studied arbuscular mycorrhizal fungus (AMF), comprises approximately 50% repetitive sequences that include transposable elements. Despite their abundance, two-thirds of TEs remain unclassified, and their regulation among AMF life-stages remains unknown. Here, we aimed to improve our understanding of TE diversity and regulation in this model species by curating repeat datasets obtained from chromosome-level assemblies and by investigating their expression across multiple conditions. Our analyses uncovered new TE superfamilies and families in this model symbiont and revealed significant differences in how these sequences evolve both within and between *R. irregularis* strains. With this curated TE annotation, we also detected that the number of upregulated TE families in colonized roots is four times higher than in the extraradical mycelium, and their overall expression differs depending on the plant host. This work provides a fine-scale view of TE diversity and evolution in model plant symbionts and highlights their transcriptional dynamism and specificity during host-microbe interactions. We also provide Hidden Markov Model profiles of TE domains for future manual curation of uncharacterized sequences (https://github.com/jordana-olive/TE-manual-curation/tree/main).

## Introduction

Arbuscular mycorrhizal fungi (AMF) are an ancient group of plant symbionts capable of colonizing thousands of different species. The AMF provides vitamins and minerals from the soil to the plants and receives carbohydrates and lipids in return ^1^. This relationship may have been established since plants first conquered the land ^2^, and there is evidence showing that AMF improve plant growth and ecosystem productivity ^3^. More recently, studies reported that these symbionts can retain atmospheric carbon, indicating they play a significant role in global carbon sequestration ^4,5^. Nonetheless, despite their relevance for plant fitness and long-term evolution, AMF show low morphological variability with no apparent plant specificity ^6^.

In addition to being ecologically and agriculturally important, AMF harbour peculiar cellular features. Their spores and hyphae always carry thousands of nuclei, and to date, no stage where one or two nuclei co-exist in one cell has ever been observed ^7^. Furthermore, despite their longevity, no sexual reproduction has been formally observed in these organisms. However, evidence has now shown that AMF strains can either carry one (AMF homokaryons) or two nuclear (AMF heterokaryons) genotypes in their cells, a genetic characteristic found in sexually multinucleated strains of ascomycetes and basidiomycetes, suggesting that sexual reproduction does exist in these prominent symbionts ^8,9^.

The low morphological diversity of AMF masks remarkable differences in gene content and structure in this group. For example, species vary in genome size and gene counts ^10^, and the variation in genome size is significantly correlated with the abundance of transposable elements (TEs) ^11^. Recent studies based on chromosome-level assemblies of *Rhizophagus irregularis*, also showed this model species carries two different (A/B) compartments, which are reminiscent of the “two-speed genome” structure previously reported in plant pathogens ^9,12^. Overall, the A-compartment is gene-rich and mostly composed of euchromatin, whereas the B-compartment contains high concentration of TEs and is mainly composed of heterochromatin ^13^. Accordingly, in AMF, the A-compartment carries most “core genes” – i.e. genes shared by all members of the same species - and shows significantly higher gene expression, while the B-compartment has a higher density in repetitive DNA, as well as in secreted proteins and candidate effectors involved in the molecular dialogues between the partners of the mycorrhizal symbiosis ^12^.

TEs are repetitive DNA classified into retrotransposons (Class I), which use RNA molecules as intermediate for “copy-and-paste”, and DNA transposons (Class II), which spread through “cutting-and-pasting” the DNA ^14^. Each class is divided into orders and superfamilies based on pathways of transposition and phylogenetic relationships ^15,16^, and thus classifying TEs is important to describe the evolution of the genome and infer its impact on the biology of any organism ^17,18^. For example, by modifying chromatin status and attracting transcription factors, these elements can also promote regulation of gene expression ^19^.

In the model AMF *R. irregularis*, about 50% of the genome is composed of TEs ^9,12,20^. Their higher abundance in the B-compartment is linked to higher rates of rearrangements ^9^, and it was proposed that elevated TE expression in germinating spores may lead to new expansions of TEs in this AMF species ^21^. Despite recent findings, key questions regarding the diversity and evolution of TEs, as well as their role in mycorrhizal interactions, remain unanswered. For example, approximately two-thirds of TEs remain unclassified, making it difficult to infer their function in AMF genome biology and evolution ^21,22^. Similarly, because analyses of TE expression have so far centered on germinating spores, it is unknown how these elements are controlled during host colonisation, and whether some show host-specific regulation. The present study addresses these questions by providing an improved classification of TE families in all *R. irregularis* strains with chromosome-level assemblies and by investigating their expression among multiple hosts.

## Material and methods

### Curation and classification of TE families

We used chromosome level assemblies of five homokaryotic (4401, A1, B3, C2 and DAOM197198) ^12^ and four heterokaryotic strains (A4, A5, G1 and SL1) ^9^ **Supplementary material S1**) as a source to build repeat libraries. The curation for nonmodel species followed the most recommended guides ^17,23^. Firstly, the repeat libraries were generated using RepeatModeler2.0.3 ^24^ with -LTRstruct mode for detecting Long Terminal Repeats sequences implemented by LTRharvest and LTR_retriever. The libraries from all strains were merged to create a single reference for the curation.

The unique library was submitted to TEclass which separates the sequences into the order level: nonLTR, LTR or DNA ^25^. This step helped us to distinguish orders with similar protein domains, such as DIRS (nonLTR) and Crypton (DNA). Tirvish, from genome-tools tools (http://genometools.org/), was used to detect Terminal Inverted Repeats (TIR) in DNA transposons elements (for order DNA/TIR) with the parameter - mintirlength 8. Hidden Markov Model (hmm) profiles of specific TE superfamilies or orders domains were generated from a combination of conserved regions described in **Supplementary material S2**. The sequences for each domain were first aligned using MAFFT ^26^, converted to stockholm format using esl-reformat and finally submitted to hmmbuild (version 3.1b2) to generate the hmm profiles ^27^.

Lastly, we provided hmm profiles for detecting elements with reverse transcriptase (LINE, DIRS, PLE, LTR, Bel, Copia, Gypsy), specific transposase superfamilies (Academ, CMC, Ginger, KDZ, Kolobok, MULE, Merlin, Novsib, P, PIF-Harbinger, PiggyBac, Plavaka, Sola-1, Sola-2, Sola-3, Tc-Mariner, Transib, Zator and hAT) and tyrosine recombinase (DIRS and Crypton) (https://github.com/jordana-olive/TE-manual-curation/tree/main/TE-domains). The open reading frames (ORF) from the reference library were generated using the tool getorf with 200 amino acids as minimum size (https://www.bioinformatics.nl/cgi-bin/emboss/help/getorf). The hmmrsearch (version 3.1b2) was used to find the sequences with TE domains using the abovementioned procedure, and by selecting the best scores using HmmPy.py (https://github.com/EnzoAndree/HmmPy). Approximately 8% of *R. irregularis* genomes consist of high-copy-number genes, known as expanded genes (e.g. Sel1, BTB, Kelch, Protein Kinase, TPR). These expanded genes may also harbor (partial) TEs insertions ^21^, and were thus removed from downstream analyses to avoid biases due to the chimeric nature of their TEs. The models can be accessed at https://github.com/jordana-olive/TE-manual-curation/tree/main/expanded-genes.

Class II elements (DNA/TIRS) were retained in the final library based on the following criteria: identified as DNA by TEclass, had a match with transposon domain from hmmsearch, harbored a TIR sequence, and size ranging between 1kb to 17kb.

Sequences with TIRS, ranging between 50bp to 1kb and lacking transposase domains were classified as MITEs. Following the hmmrsearch, certain elements exhibited similarity with more than one domain from different orders due to close relationships. To achieve the most accurate classification, these sequences were analyzed using phylogenetics to determine their evolutionary relationships. For this work, the protein sequences corresponding to these elements were aligned using MAFFT ^26^, and submitted to RAxML ^28^ to produce a phylogenetic tree using the PROTGAMMA model with 1000 bootstraps. The best tree resolution was visualized using ggtree in R ^29^. The final classification of sequences aligning with more than one domain was based on their relationships, clustering elements from the same orders together, as illustrated in **Supplementary Figure S1**.

Throughout the curation process, TEs sequences classified by RepeatModeler were retained only if accurately identified by the program TEclass ^25^ and if the respective transposition domain was identified within their sequence. For newly identified sequences that were previously labelled as “unknown” by RepeatModeler, the classification was determined based on TEclass ^25^, the presence of a transposition domain, and their relationship to known sequences based on the phylogenetic reconstruction. The final library, and models are available on https://github.com/jordana-olive/TE-manual-curation/tree/main and can be applied in any other dataset to custom TE characterization.

### Repeat landscapes of genomes and compartments

We ran the RepeatMasker (version 4.1.2-p1) ^30^ for all strains, with the parameters -a -s, using the curated library as the reference (-lib option). The repeat landscapes were generated from modified createlandscape.pl and calculedivergence.pl scripts, provided in RepeatMasker files. These scripts calculate the divergence levels between the alignment of each TE sequence and the consensus family in the reference library. The landscapes also were generated to A and B-compartment, which are currently available only for the strains DAOM197198, A1, C2, A4 and A5 ^9,12^. To assess whether the distribution of TEs varies across compartments, we extracted the percentage values from the landscapes of each Kimura bin, and then conducted a paired t-test using R. A significant difference between the landscapes was considered when p < 0.05.

### Transposable element and gene expression analysis

We evaluate the expression of genes and TEs using available RNAseq from different tissues (germinating spores ^21^, intraradical mycelium, arbuscules ^31^ and extraradical mycelium ^32^), and mycorrhized roots from different plant hosts colonized by DAOM197198 (*Allium schoenoprasum, Medicago truncatula, Nicotiana benthamiana* ^31^ and *Brachypodium distachyon* ^33^). The accession numbers to the data are available in **Supplementary material S3**. The reads were filtered using Trimmomatic ^34^ and aligned to DAOM197198 reference genome using Bowtie2 ^35^. The read count was accessed by TEtranscripts ^36^ guided by the TE annotation performed in this study and gene annotation executed by ^12^. Using DESeq2 ^37^, for each host condition and tissue, the differential expression was generated comparing germinated spores as control. A transcript was considered differentially expressed when padj (adjusted p-value) is equal or lower than 0.05.

### Transposable element nearby gene correlation

TE location and gene expression correlation was analysed using available RNAseq from DAOM197198 generated through Oxford Nanopore Technology (ONT) sequencing ^20^. The long ONT-RNAseq reads were filtered and trimmed using pychopper (https://github.com/epi2me-labs/pychopper) (7.98% did not pass the quality parameters and were discarded). The nucleotide correction was performed based on self-clustering using isONTcorrect ^38^. The filtered and corrected reads were aligned to DAOM197198 genome using hisat2 and annotated using stringtie ^39^ guided by TE annotation performed in this study and gene annotation executed by ^12^. In the same way, stringtie generated the counts of the transcripts, used in the expression analysis. For detecting TEs upstream of genes, we extracted the genomic regions up to -1000 to the transcript start position and then intersected with TE annotation using bedtools ^40^. The Pearson correlation method in R was employed to assess the co-expression between genes and their upstream TE pairs.

## Results

### A curated database reveals new TE families in Rhizophagus irregularis

Using RepeatModeler and RepeatMasker, recent analyses of *R. irregularis* chromosome-level datasets indicated that strains of this species carry an average of 50% of TEs ^9,12^. However, only one third of their repeat content could be classified, and thus on average 30% of all available genomes are composed by unclassified TE sequences ^9,12,20,22^. To address this, we used chromosome-level assemblies from 5 homokaryon and four heterokaryon *R. irregularis* strains to generate curated repeat libraries. When all genomes are considered, out of total of 9,257 TEs sequences identified by RepeatModeler, only 2,369 (approx. 25%) can be considered well defined, *bona-fide* non-redundant consensus sequences harboring transposition domains. The notable reduction in TE numbers is due to non-curated datasets containing highly degenerated TE without domains (relics), and repeats that cannot be classified with current knowledge of TE evolution.

In the final curated library, 1,458 sequences belong to families previously classified by RepeatModeler (K, **Figure 1a**), while 636 sequences represent newly identified families (N, **Figure 1a**) of the following orders; SINE, LINE, LTR, DNA/TIRS, DNA/Crypton, Maverick and RC/Helitron (**Figure 1a**). Following curation, in the DAOM197198 genome, sequences belonging to these orders increased in number by 4-fold on average, and similar results were obtained for all investigated *R. irregularis* strains (**Figure 1b**). We also assessed the influence of the manually curated library on the repeat landscape of the DAOM197198 genome (see **Figures 1c-e**). Following TE curation, the percentage of the DAOM197198 genome is represented by classified TE increased from 12% (**Figure 1c-d**) to 36% (**Figure 1e**).

**Figure 1.**
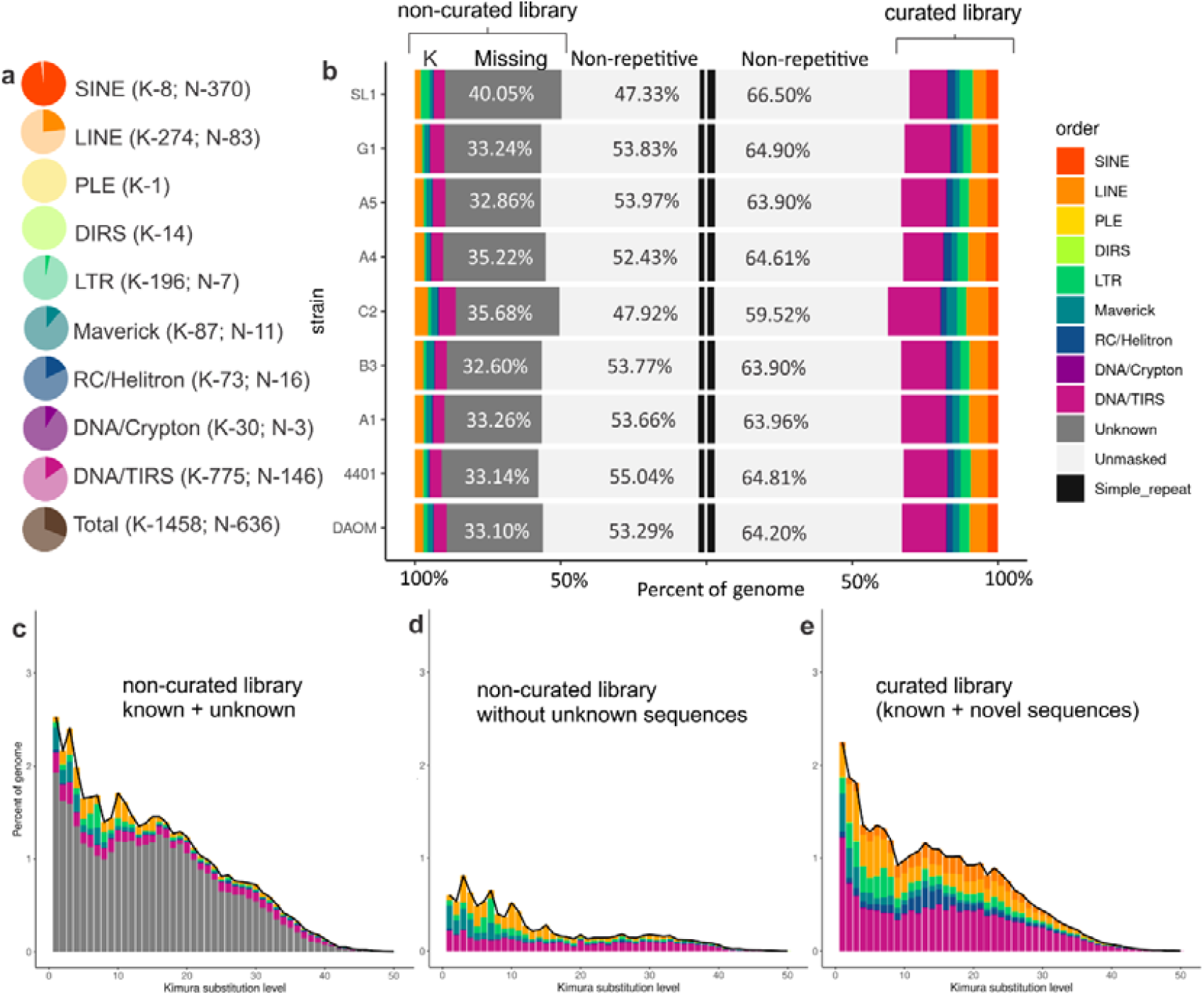
Description of the curated TE library. **a** Proportion of known (detected by RepearModeler, in light color, K) and novel (in bold colors, N) curated sequences. **b** RepeatModeler and curated library annotation comparison across the strains. The *y-axis* represents the *R. irregularis* strains, while the *x-axis* shows the relative abundance of the TEs in the genome from the homology-based annotation (RepeatModeler, left) and the current annotation (curation, right). **c-e** Repeat landscape of DAOM197198 using different libraries as a source for annotation. Each plot shows the sequence divergence from its consensus (*x-axis*) in relation to the number of copies in the genome (*y-axis*). Newer insertions are shown in the left peaks, while the older and more degenerate insertions are on the right side of the graph. **c** Annotation performed using the non-curated library from RepeatModeler, which depicts the unknown elements (in gray). **d** Only non-curated TEs detected by RepeatModeler, which comprehends 12% of the repeat content. **e** Landscape generated with the final curated library.

The newly curated *R. irregularis* TE library now includes novel families of non-LTR retrotransposons: SINE, LINE/CR1-Zenon, LINE/L1-Tx1, LINE/L1-R1, LINE/R2-Hero, LINE/RTE-BovB, LINE/I, which were detected based on phylogenetic analyses (**Supplementary Figure S1**). A novel superfamily of DNA transposons – e.g., Transib, - and novel families belonging to known AMF TE superfamilies were also detected, including Crypton, CMC, hAT-19, MULE, Sola-2, Tc1-Mariner and Plavaka (**Figure 2**). The Plavaka family, which is part of CACTA/CMC/EnSpm superfamily, is particularly prevalent in DAOM197198, A1, B3, C2, A5 and G1. This family has already been identified in fungi ^41,42^, but is not deposited in publicly available repeat databases, which is why the RepeatModeler could not detect this family.

**Figure 2.**
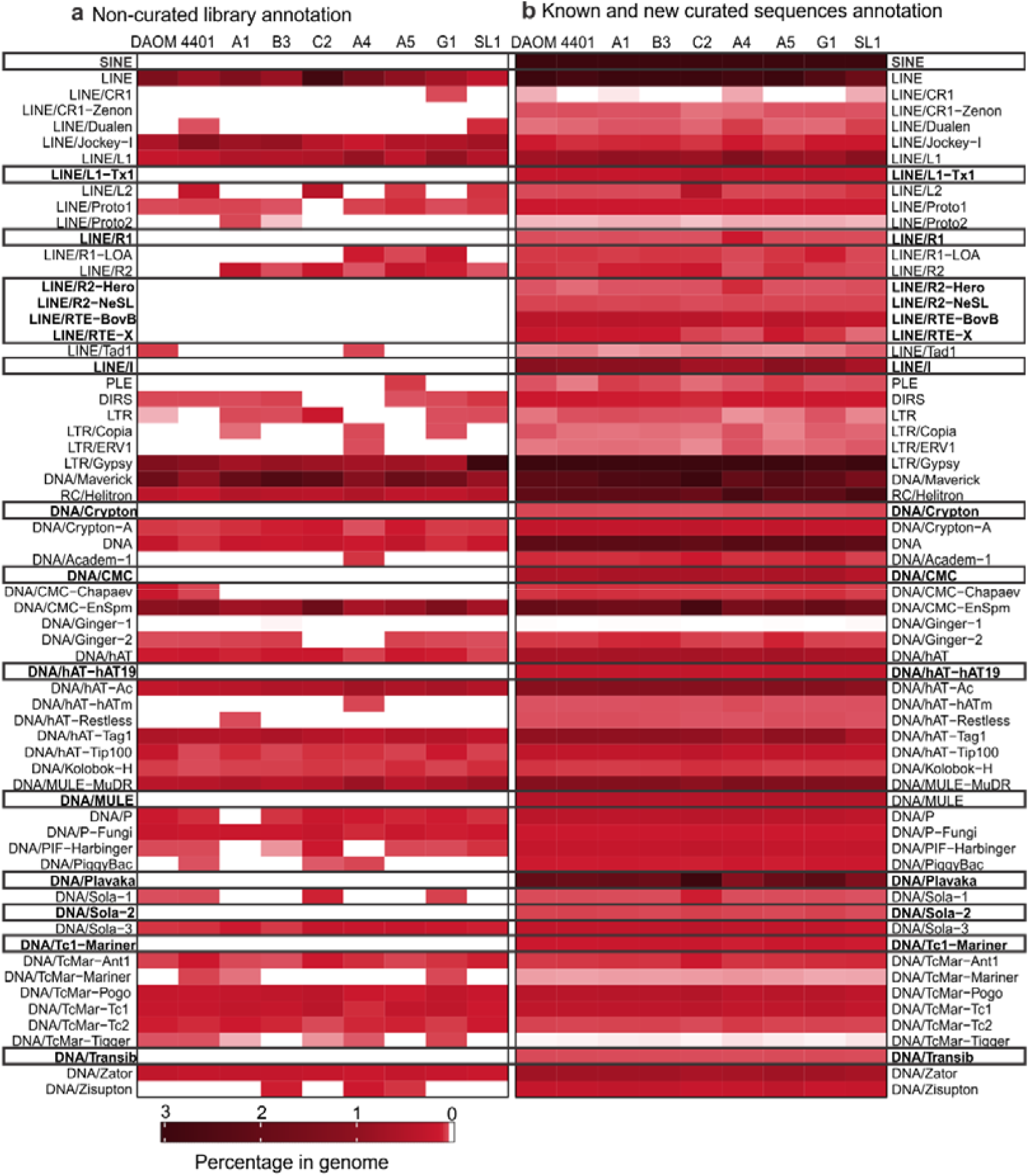
Annotation of TE families in *R. irregularis* strains. The heatmaps show the percentage of families in each strain considering the RepeatMasker annotation. **a** Percentage of TE families in each genome using the non-curated library. **b** Annotation using the curated library with known and novel TE families. Blank areas mean absence of certain family in that annotation, while the black rectangles highlight the new families found in the curation.

### Repeat landscapes differ among and within R. irregularis strains

In the *R. irregularis* isolate DAOM197198, TEs are mobilized and retained differently across the genome ^21^, with two main waves of expansions. We accessed the distribution of TEs in the other strains based on Kimura substitution calculation, which estimates the level of divergence of annotated TE compared to its consensus in the reference library. A lower number of Kimura substitution indicates high similarity between annotated sequences and their consensus, suggesting recent insertion events, while a higher number indicates greater evolutionary distance and likely older TE insertions.

Our analysis revealed that the patterns of repeat landscapes based on Kimura substitution levels differ among *R. irregularis* strains. Specifically, in most strains, expansions of TEs are recent, as highlighted by the high number of elements with Kimura substitutions ranging from 0 to 5, supporting the findings that new TEs expansion bursts exceed older TE insertions in DAOM197198 ^21^ (**Figure 3**). However, other strains exhibit different TEs distributions. For example, strains A4 and SL1 carry a larger proportion of TEs with older Kimura substitution levels, indicating strain-specific patterns of TE emergence, evolution, and retention. Remarkably, the genome of C2 has a much larger proportion of younger, and thus likely more active TEs, and it is plausible that this feature is linked to the larger genome size of this strain compared to relatives - i.e., the C2 genome is 162Mb compared to an average of 147Mb for other *R. irregularis* strains (**Figure 3**).

**Figure 3.**
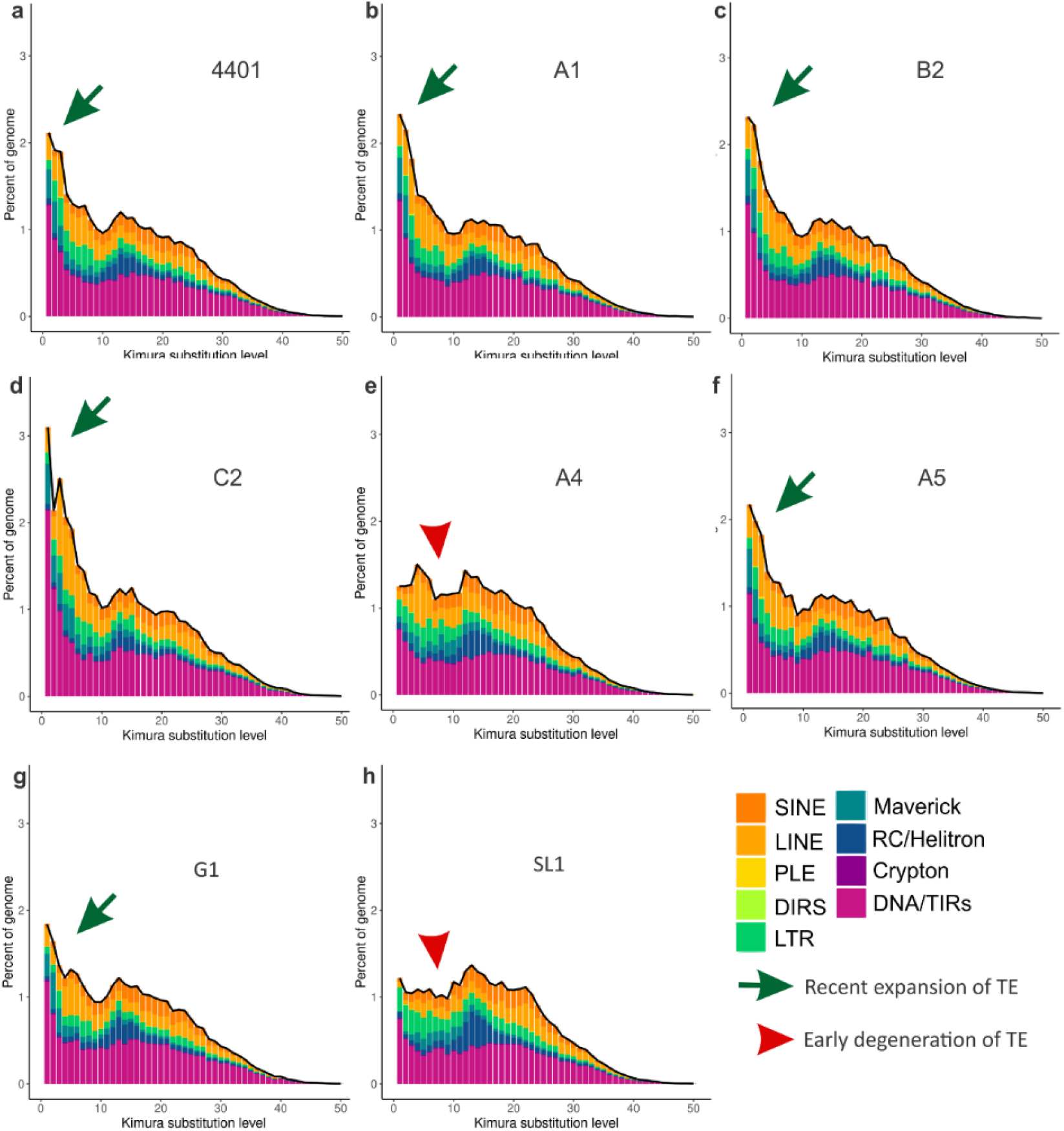
Repeat landscape of *R. irregularis* strains. **a-h** Each plot shows the sequence divergence from its consensus (*x-axis*) in relation to the number of copies in the genome (*y-axis*). Newer insertions are shown in the left peaks, while the older and more degenerate insertions are on the right side of the graph. The strains with recent burst of TE expansions are indicated with the green arrow, while the ones representing early TE degeneration are highlighted with red arrow.

The A and B-compartment were recently identified in the chromosome-level assembly of three homokaryons (DAOM197198, A1 and C2)^12^ and two heterokaryons (A4 and A5)^9^ strains. Based on available data, we observed that the distribution of TEs based on Kimura substitution levels also differ significantly between A and B-compartment in all strains (**Figure 4**). Specifically, the proportion of TEs with Kimura substitutions between 10 and 20 – i.e., older TEs - is always higher in the B-compartment (p<0.05), indicating that these elements are maintained over time at higher levels in this portion of the genome. In contrast, old and degenerated TEs appear to be rapidly eliminated in the A-compartment (**Figure 4**).

**Figure 4.**
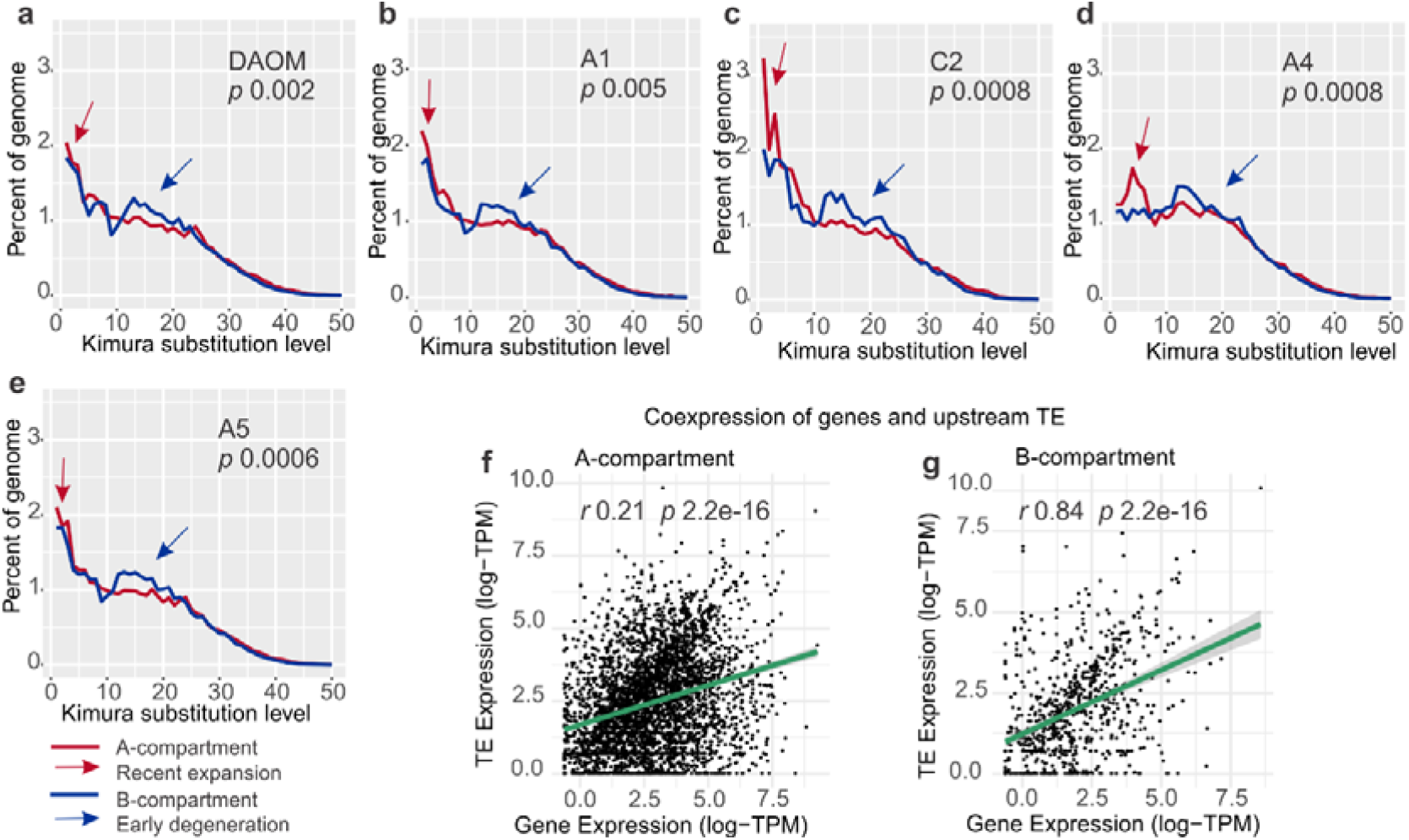
**a-e** Repeat landscapes in *R. irregularis* strains with annotated A and B-compartments. The arrows point to the divergence between the compartments. **f-g** Coding genes and upstream TEs correlation of expression in DAOM197198. Each point represents a coding gene-upstream TE pair, where the *y-axis* is the TE expression (logTPM) and the *x-axis* is the coding gene expression (logTPM). The green line represents the regression, followed by the confidence interval in light grey.

### TE and gene expression is positively correlated within the B-compartment

The localization of TEs nearby genes can impact their expression ^19,43^. We investigated whether the presence of TEs upstream of genes is associated with gene expression in compartments by identifying the expression of TEs located within 1000 nucleotides upstream of genes using long-read RNA data from DAOM197198 strain ^20^. We found no correlation between the expression of TEs and coding genes located in A-compartment (*r*=0.21, p<0.0001) (**Figures 4f**), however, a positive and significant correlation exists for genes and TEs in the B-compartment (*r*=0.84, p<0.0001) (**Figures 4g**). This finding suggests that TEs and genes in B-compartment are being co-expressed.

### Transposable elements are significantly more expressed in colonized roots compared to extraradical mycelium

To obtain additional insights into the biology of TEs, we investigated their regulation during host colonization using *R. irregularis* DAOM197198 RNA-seq data from multiple tissues, including micro-dissected cells of arbuscules (ARB) and intraradical mycelium (IRM), and extraradical mycelium (ERM) (**Figure 5a**) in symbiosis with *Medicago truncatula* roots (**Figures 5b-d**).

**Figure 5.**
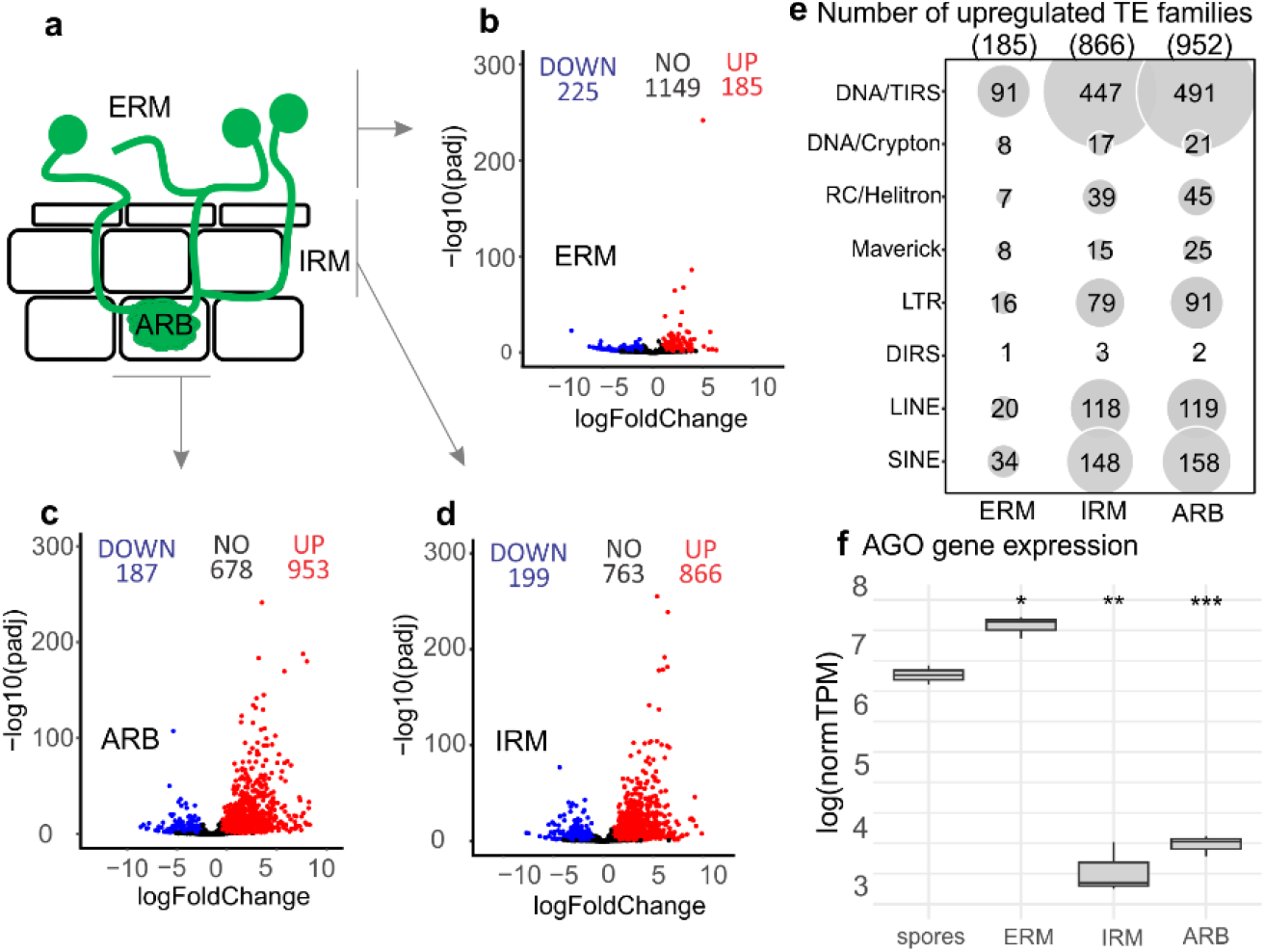
Expression of TEs in *R. irregularis* through life-stages of colonization. **a** Schematic view of the different life stages compared to spores. ERM, extraradical mycelium; IRM, intraradical mycelium, ARB; arbuscules. **b-d** Volcano plots of expressed TEs in each colonization stage, evidencing the number of down and upregulated sequences, in blue and red, respectively. The ones expressed but not significantly different from spores are shown in black as NO. **e** Number of upregulated TEs in each condition. Inside the parentheses are the number of total upregulated TEs in that specific condition. The gray circle size is according to the number of upregulated sequences. **f** *AGO-like* gene (Rhiir2092) expression under different conditions. Colonization stages collected by laser dissection of ARB and IRM; ERM and mycorrhized roots. The t-test compared all the conditions with spores, where: * p < 0.05; ** p < 0.001 and ns is no significance. The box limits represent the range of 50% of the data, the central line marks the median, and the vertical lines out of the boxes are the upper and lower values. normTPM – normalized transcripts per million.

The number of upregulated TEs is more than four times higher in IRM and ARB than in extraradical mycelium in the same host (**Figure 5b-d**). TEs significantly upregulated in colonized tissue (ARB and IRM) include DNA/TIRS, LTR, LINE and SINES orders. Among these, LTR/Gypsy, LINE, CMC-EnSpm, Plavaka, hAT-Tag1, hAT-Ac and Helitron are the superfamilies with more expressed families (**Supplementary Material S4**).

One mechanism to control TEs mobilization is through RNAi and AGO proteins ^44^. Typically, fungi possess 1-4 AGO genes per genome ^45^. However, the *R. irregularis* harbors 40 copies of *AGO-like* genes, and of these 25 contain all typical AGO core domains (e.g. PIWI, PAZ, MID and N-terminal) with some exhibiting signs of expression ^46^. Among these *AGO-like* genes, only one (Rhiir2092, JGI 1582012) presents significant expression – i.e. more than >100 transcripts per million (TPM) in the samples, while the remainder exhibit minimal to no expression. We find that the Rhiir2092 gene is significantly more expressed in ERM (p < 0.05) compared to the other conditions (**Figure 5b** and **f**). Specifically, for this gene, arbuscules and intraradical mycelium laser dissected cells have significantly reduced expression, which again differs from the significantly higher expression of TEs in these conditions (p < 0.05) (**Figure 5b** and **f**).

### TE regulation changes with different hosts and correlates with host genome size

Analyses of RNA-seq data roots from multiple hosts colonized by DAOM197198 (*Allium schoenoprasum, Medicago truncatula, Nicotiana benthamiana* and *Brachypodium distachyon*), also reveals that TE expression differs significantly among hosts, with some families being expressed only in one of four hosts (**Figure 6** and **Supplementary Material S4**). For example, a total of 404 families are upregulated during colonization with *A. schoenoprasum* roots (**Figure 6a**), including both families from Class II elements (e.g., Sola-1, hATm, Academ-1, DIRS) and Class I elements (e.g. CR1-Zenon and DIRS) that are only upregulated with this host. In *N. benthamiana* and *M. truncatula* roots, the symbiont upregulates a smaller number of TE families compared to *A. schoenoprasum*, 251 and 197 respectively, while *B. distachyon* was the condition with lowest TE upregulation overall (116 families) (**Figure 6d**). The differences in TE expression among hosts are not linked to the variability in expression of AMF *AGO-like* gene – i.e., hosts with highest TE expression do not always have low *AGO-like* expression and vice-versa.

**Figure 6.**
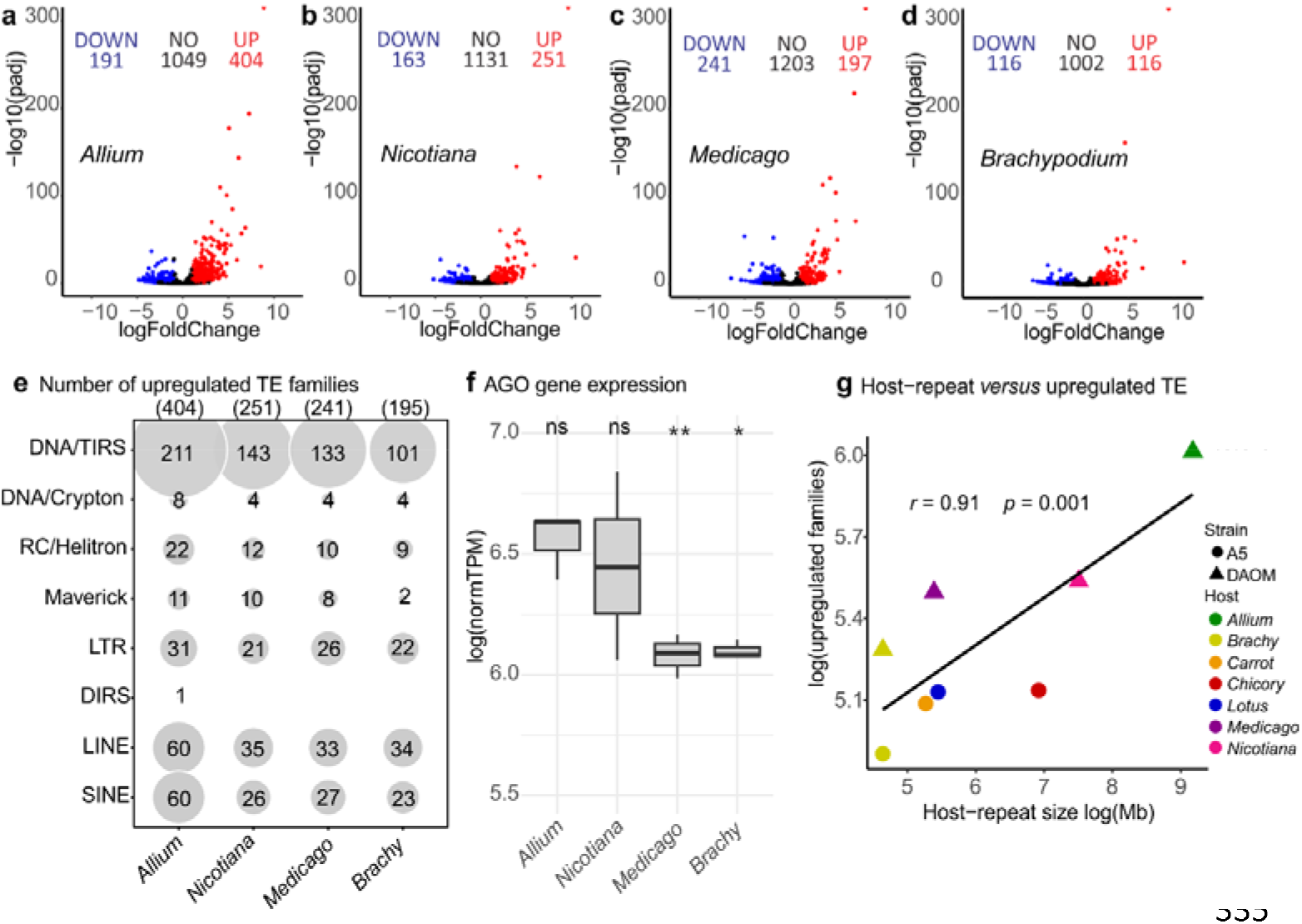
Differential expression of TEs in DA OM 197 198 duri ng symbiosis with different plant hosts compared to germinated spores. **a-d** Volcano plots showing the down and upregulated TEs in relation to spores, in blue and red respectively, considering padj <0.05. In black are the ones with no significant difference in expression in relation to the control. **e** Number of overexpressed TE families in each host. **f** *AGO-like* gene (Rhiir2092) expression under colonization with different hosts. The t-test was performed comparing all the conditions with spores, where: * p < 0.05, * p < 0.001 and ns is no significance. The box limits represent the range of 50% of the data, the central line marks the median, and the vertical lines out of the boxes are the upper and lower values. normTPM – normalized transcripts per million. **g** Correlation between the repeat content of the host genome (in base pairs) and the number of families overexpressed in the AMF during the symbiosis. The repeat size of the hosts was calculated based on the most recent genome assembly and annotation: *Allium sp*.^48^, *Brachy*^49^, Carrot^50^, Chicory^51^, *Lotus japonicus*^52^, *Medicago truncatula* ^53^, *Nicotiana benthamiana* ^54^. Brachy – *Brachypodium*.

Given the observed variation in TE expression in the AMF, we wondered how the plant host influences the differential expression of these superfamilies. A significant and positive correlation (*r =* 0.91, *p* = 0.001) between the repeat content of the host genome and the number of overexpressed families in the symbiont. For example, *A. schoenoprasum* (14Gb) is the host with the most TE content and its colonized roots have the highest number of TEs upregulated in the AMF. In contrast, *B. distachyon* (272Mb) has the lowest TE content and has significantly lower TE upregulation in the symbiont.

It is noteworthy that, in contrast to fungi, the repeat content of plant species has been well characterized using available tools, because most repeat databases have been curated using plant genome information ^24,47^. As such, it is very unlikely that the significant correlation we observed were skewed by the non-curated nature of the plant genomes we used, particularly given that these findings are consistent among multiple plant hosts and conditions.

## Discussion

### An improved view of TE family diversity and evolution in a model plant symbiont

Through curation of *R. irregularis* repeat libraries, we first improved the proportion of annotated families in these model plant symbionts and uncovered novel sequences representing the largest proportion of their repetitive sequences. Our work also revealed how *R. irregularis* strains differ in TEs retention and deletion over time within two genome compartments. For example, our findings indicate that strains with largest genome sizes (C2) show a combination of a higher rate of TEs emergence and retention of these elements. We also uncovered notable differences in how TEs evolve within each strain, with some (A1, B3) having have much higher proportions of very young TEs compared to others (SL1, A4) that carry levels of Kimura substitution rates indicative of early repeat degeneration and fewer recent expansion.

### TE retention rates are different between compartments

By investigating the degree at which TEs accumulate mutations over time, a significant distinction emerged between A/B compartments. Specifically, all A-compartment landscapes show a continuing invasion of novel/young TE insertions, and the high methylation present in this compartment ^12^, and/or purifying selection, might be needed for their control and rapid removal; as evidenced by the lower TE density in this compartment. In contrast, these insertions accumulate in the B-compartment, leading to a notable inflation of these elements over time, as shown by the stable TE density along the axis that defines the Kimura substitution rates. The accumulation of TEs in the B-compartment might be linked with their domestication ^55,56^ and/or the emergence of new functions; a view supported by the significant positive correlations we observed in expression of TEs and genes and by similar findings from plant pathogens ^57,58^.

### TE regulation and evolution further underpin similarities between AMF and known fungal pathogens

Obvious similarities between the genomes of AMF and those of plant pathogens have been known for some time ^59,60^. These include enrichments in transposable elements, and genomes subdividing into highly diverging regions dense in effector genes and TEs, and more conserved ones composed of core genes and low repeat density.

In the plant pathogen *Verticilium dahliae*, TEs often locate in proximity to highly expressed pathogenicity-related genes within fast evolving adaptive genomic regions ^61^. These regions are reminiscent of *R. irregularis* B-compartments ^12,60^, which are also enriched in TEs and secreted proteins that promote symbiosis with different hosts ^62^. As such, the significant correlation we observed between TEs and genes specific to the B-compartment may mirror identical processes in AMF and plant pathogens.

The increased upregulation of genes in close proximity to TEs during the colonization stages in the B-compartment could also indicates a TE control for de-repressing those regions of the genome ^63^. Indeed, it has been proposed that TE-effectors regulation is well-timed; i.e. both are expressed during the infection and repressed in the absence of the host ^58^. In filamentous fungi, the variability of TEs can also allow for escaping mechanisms of recognition by the plant immunity system ^57,58^, and it is thus possible that in AMF the expression of transposable elements could aid plant-symbiont communication during colonization.

### What drives TE upregulation and host-specificity during colonization?

TE expression is active in germinating spores ^21^, but our study shows their expression in colonized roots is much higher still. In this context, we observed that an *AGO-like* gene exhibits significantly higher expression in extraradical mycelium compared to colonized roots ^46^. AGO proteins are known TE regulators^64^, and thus our results suggest that RNAi is one of the key factors implicated in the regulation of TEs across stages of the mycorrhizal symbiosis -i.e. downregulation in extraradical mycelium, and upregulation *in planta*. In fungi, it is expected 1-4 AGO proteins to regulate TEs ^45,65^. However, the genome of *R. irregularis* unexpectedly contains an expanded set of 40 copies of this gene ^46^. The remaining *AGO-like* genes, which exhibit minimal or no expression, may be implicated in other functions ^46^. This could be attributed to these copies being incomplete or carrying additional domains ^46^. Notably, the expression of this *AGO-like* gene did not vary significantly among hosts, and thus other factors could be responsible for the host-specific variation in the TE expression that we observed.

One intriguing possibility is that TE expression in the hosts directly or indirectly influences TE expression in the AMF, as seen in multiple plant-microbe interactions in cross-talk regulations ^66–69^. In support of this view, we found a positive correlation between TE abundance in the host and TE expression in AMF. With mounting evidence of molecular cross-communication between the mycorrhizal partners, including RNA from the fungi interacting with mRNA from the hosts ^46,70,71^, it is likely that molecular dialogues between mycorrhizal partners also result in the increased TE expression we observed in the fungal symbiont.

## Acknowledgements

Our research is funded by the Discovery program of the Natural Sciences and Engineering Research Council (RGPIN2020-05643) and a Discovery Accelerator Supplements Program (RGPAS-2020-00033). NC is a University of Ottawa Research Chair in Microbial Genomics. JINO is founded by Mitacs Accelerate Program (IT16902) and a Discovery Accelerator Supplements Program (RGPAS-2020-00033).

## Author contributions

JINO and NC designed the study. JINO performed the experiments and analysis of the data. JINO and NC wrote the paper. NC mentored and supervised all the processes.

## Data availability

The genomes and RNAseq used in this study are described in **Supplementary material S1** and **S3**. The ONT RNAseq can be accessed at SRR21968700.

**Supporting information**

**Supplementary file S1 –** Chromosome level assemblies of *R. irregularis* strains used in this study.

**Supplementary file S2 –** Domains used to construct the hmm models.

**Supplementary file S3 –** RNAseq data and alignments stats used in this study.

**Supplementary file S4 –** Number of upregulated TEs in each condition.

**Supplementary figure S1 –** Phylogenetic tree of TEs from different orders with similar domains. Figure in high resolution can be downloaded here https://github.com/jordana-olive/TE-manual-curation/blob/main/Supplementary-figure-S1.png.

