## Supplementary Figure S1 for "Strain-specific evolution and host-specific regulation of transposable elements in the model plant symbiont *Rhizophagus irregularis*"

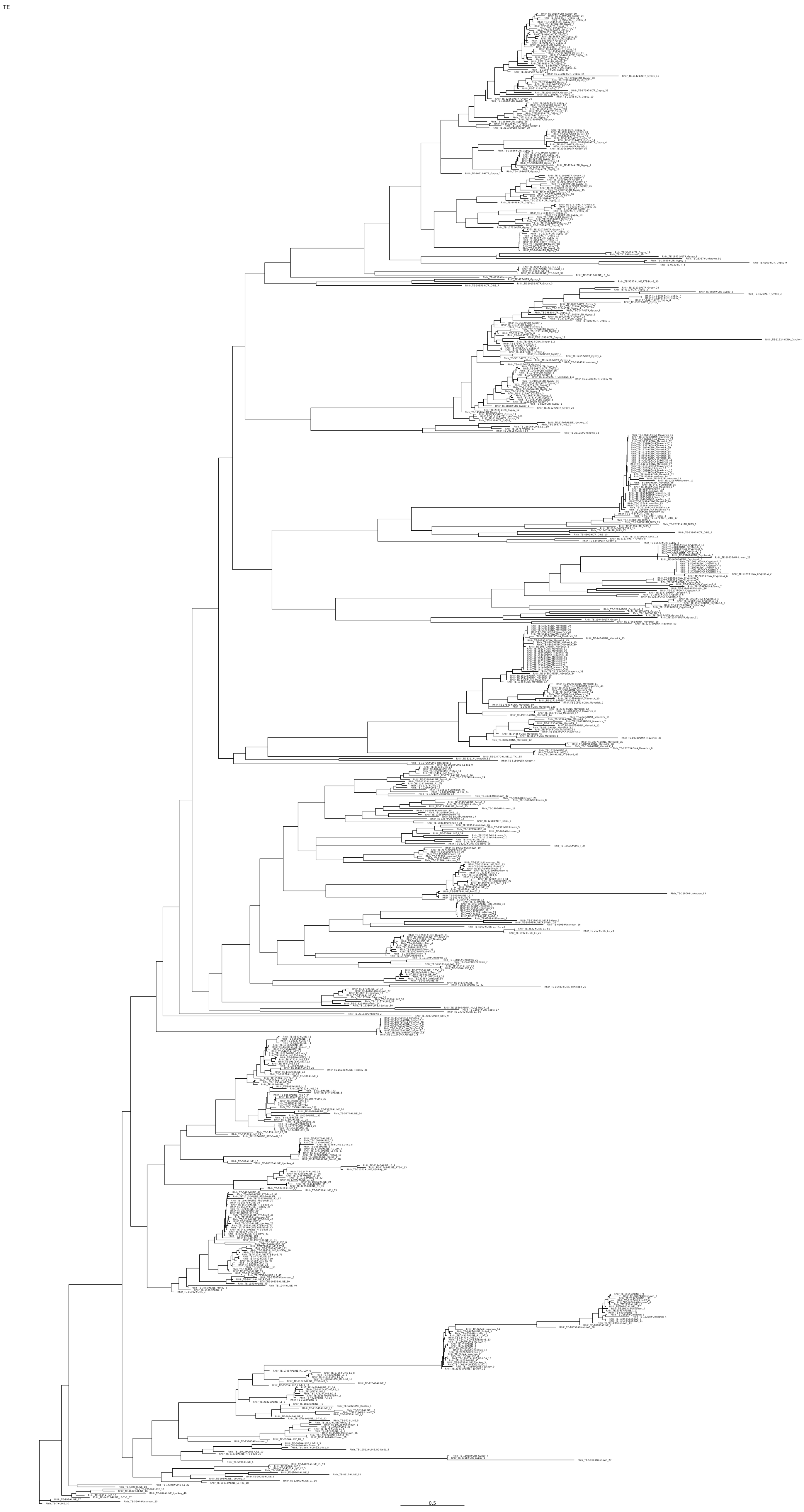


**Supplementary figure S1** - Phylogenetic tree of TE from different orders with similar domains, [access it in high quality]
